# Leptin drives osteoarthritis pain through sensory neuron reprogramming independent of cartilage signaling

**DOI:** 10.64898/2026.05.22.727205

**Authors:** Hope D. Welhaven, Elise K. Truchan, Kristin L. Lenz, Bethany A. Andoko, Michael R. Mazzucco, Reyna E. Villa, Arin K. Oestreich, Bo Zhang, Dana E. Orange, Joseph B. Lesnak, Theodore J. Price, Farshid Guilak, Kelsey H. Collins

## Abstract

**Objective:** Pain in osteoarthritis (OA) is often discordant with structural joint damage, particularly in obesity-associated OA, where adipose-derived signals may drive nociception independently of cartilage pathology. Leptin has been demonstrated to be necessary, but not sufficient, to drive obesity-associated OA. Here, we tested the hypothesis that leptin mediates OA-associated pain through sensory neuron reprogramming rather than chondrocyte-intrinsic signaling, suggesting a fat-sensory nerve axis.

**Design:** Male and female constitutive leptin-deficient (Ob/Ob), heterozygous (Ob/+), and wild-type (WT) mice, as well as chondrocyte-specific leptin receptor knockout mice (Aggrecan-CreERT2;LepRfl/fl), were challenged with destabilization of the medial meniscus (DMM) surgery to induce OA. Pain-related behaviors, joint pathology, serum cytokines, and lumbar dorsal root ganglia (DRG) transcriptomes were assessed. Human DRG cultures treated with leptin underwent transcriptomic profiling. Secondary analyses of human infrapatellar fat pad and synovium single-cell datasets evaluated leptin and leptin receptor expression patterns.

**Results:** Chondrocyte-specific deletion of the leptin receptor did not mitigate OA pathology or pain. Global leptin-deficient (Ob/Ob) mice exhibited worse structural joint outcomes than WT and Ob/+ animals following DMM yet were robustly protected from OA-associated hyperalgesia – directly dissociating pain from structural pathology and demonstrating that leptin is involved in nociceptive sensitization. Serum cytokine profiles were sex-dependent and did not align with pain outcomes, separating systemic inflammation from nociceptive differences. Transcriptomic analysis of DRGs revealed that leptin drives enrichment of lipid metabolism, eicosanoid, and inflammatory programs, whereas leptin deficiency shifts sensory neurons toward a cytoskeletal remodeling state that does not sustain pain signaling. In human DRG cultures, leptin treatment produced a transcriptomic shift to enrich for neuronal excitability while vehicle treated cells were enriched for inflammatory signaling. Human infrapatellar fat pad and synovium transcriptomic data demonstrated adipocyte-enriched leptin expression and broad distribution of the leptin receptor across stromal, vascular, immune, and adipocyte populations.

**Conclusions:** Leptin contributes to OA pain through neuro-immune crosstalk between adipose tissue and sensory neurons rather than through direct cartilage signaling. These findings identify leptin-associated neuronal programs linked to nociceptor sensitization and support targeting leptin-modulated neuro-immune pathways as a strategy to alleviate OA pain independently of structural disease progression.

## Introduction

Pain is the defining symptom of osteoarthritis (OA) and the primary driver of care-seeking behavior^1–3^, yet current pain management strategies remain inadequate and the mechanisms underlying OA-associated pain remain poorly understood. Clinically, pain correlates poorly with the degree of radiographic evidence of structural damage in OA^4^, highlighting a fundamental gap in our understanding of how joint damage is converted into perceived pain. This discordance is particularly evident in obesity, where patients frequently experience greater pain for a given level of joint degeneration compared to individuals without obesity^4^. OA prevalence is also increased in both weight-bearing and non-weight-bearing joints in obesity^3^, suggesting that factors beyond biomechanics contribute to disease pathogenesis and pain. Together, these observations support a growing view of OA as a systemic disease^5^ in which adipose-derived signals influence nociceptive outcomes independently of structural joint damage.

Leptin, an adipokine secreted proportionally to fat mass and chronically elevated in obesity, has emerged as a candidate mediator linking adiposity to OA^6–9^. Prior studies, including those from our lab^6, 7^, have demonstrated that leptin promotes cartilage damage and may be sufficient to drive cartilage damage^9^. whether leptin drives OA-associated pain through cartilage-intrinsic mechanisms or through pathways outside the joint remains unclear. Leptin receptors are expressed across multiple tissues relevant to OA – including chondrocytes, osteoblasts, osteocytes, synoviocytes, bone marrow stromal cells, and fat cells^9–13^ – indicating that leptin may exert tissue-specific effects on pain signaling. Despite this established role in structural joint damage, the downstream mechanisms through which leptin contributes to OA pain have not been defined, and therefore generating a mouse that specifically lacks the leptin receptor on chondrocytes is a tool to allow us to answer this question directly.

Critically, leptin receptors are expressed not only in joint tissues but also in sensory neurons^14^, raising the possibility that leptin acts on the peripheral nervous system to modulate nociception independent of its effects on the local joint. This distinction matters: if leptin’s contribution to pain operates through sensory neurons rather than through leptin-mediated chondrocyte activity, then the pain-structure discordance observed in obesity may have a direct molecular explanation, and therefore, a distinct therapeutic target. Sensory neurons of the dorsal root ganglia (DRG) are recognized as a critical nexus for OA pain. Knee joints are densely innervated by sensory afferents, and nociceptors in OA-affected joints become hypersensitive to stimuli, producing increased nociceptive input that drives pain directly and sensitizes central pain processing^15^. Moreover, recent work has increasingly implicated neuro-immune crosstalk outside of the joint in the development and maintenance of OA pain^16–20^. Adipokines, such as leptin, are ideally positioned to participate in this crosstalk where they are secreted systemically, reach DRGs through the circulation, and can act on sensory neuron signaling and inflammatory programs. Whether leptin directly reprograms DRG transcriptional programs to promote nociception remains to be examined.

Here, we tested the hypothesis that leptin mediates OA-associated pain through signaling between adipose tissue and sensory neurons, rather than through chondrocyte-intrinsic mechanisms, defining a fat-sensory nerve axis that links adiposity to nociception independently of joint structural damage. Using global constitutive and inducible chondrocyte-specific genetic models of leptin signaling in both male and female mice, combined with behavioral, inflammatory, and transcriptomic analyses in mouse and human DRGs, we identify leptin as a regulator of OA-associated pain that is dissociated from structural joint damage. In parallel, analysis of human infrapatellar fat pad and synovium demonstrates adipocyte-enriched leptin expression alongside broad leptin receptor distribution across multiple joint-associated cell populations. Together, these findings provide a mechanistic framework that begins to disentangle the pain–structure paradox in obesity-associated OA and identify leptin-dependent sensory neuron signaling as a potential therapeutic target for pain.

## Methods

### Animals

Chondrocyte-specific leptin knockout mice (Acan-Cre^ERT2^;LepR^fl/fl^, LepR cKO) were generated utilizing Aggrecan-Cre^ERT2^ mice (JAX #019148) and leptin receptor^fl/fl^ mice (JAX #008327). Littermate LepR^fl/fl^ mice lacking Cre were used as controls (floxed, provided by M. Myers, UMichigan). Tamoxifen was administered for 4 days at 12 weeks of age to induce Cre-mediated recombination. Global leptin knockout (JAX #000632) (Ob/Ob), heterozygous (Ob/+), and wildtype (WT) mice were used as a model of genetic obesity. All experimental procedures were approved by Washington University and the University of California San Francisco Animal Care and Use Committees. At 16 weeks of age, male and female mice underwent destabilization of the medial meniscus (DMM) to model OA in the left hindlimb, and the right hindlimb remained naive. At 28 weeks of age, 12 weeks post-DMM^6, 7, 21^, pain and behavioral testing was performed before sacrifice. Serum, synovial fluid, knee joints, and DRGs were harvested.

### Histology

Knee joints were prepared according to previous methods^6, 7, 22^. In brief, joints were fixed, decalcified, processed, and embedded prior to sectioning. Serial sections were cut (5 µm), stained with Safranin-O/Fast Green or Hematoxylin and Eosin (H&E), and evaluated using Modified Mankin, osteophyte, and synovitis scoring systems. Scoring was conducted by blinded graders and mean scores across three independent graders were used for analysis. Inter-rater reliability between scorers was high (intraclass correlation coefficient, ICC = 0.9).

### Pain and Behavioral Assessment

Pain-related and behavioral testing was blindly assessed prior to sacrifice. Tactile allodynia was assessed using Electronic Von Frey (EVF) testing. Pressure-pain thresholds of the knee were evaluated using a Small Animal ALGO meter (SMALGO, Bioseb) system^6, 22, 23^. For all behavioral assays, mice were randomized and acclimated to the testing environment and equipment prior to data collection according to established protocols^7, 22^.

### Serum Inflammatory Profiling

Fasted blood samples were collected from Ob/Ob, Ob/+, and WT mice for serum isolation. Serum cytokine and chemokine profiles were quantified using an 18-plex Luminex assay (Eve Technologies)^6, 7, 24, 25^. Leptin concentrations were measured by ELISA (R&D Quantikine).

### Dorsal Root Ganglion Bulk RNA-Seq

Bulk sequencing on L3-L5 DRGs was performed as previously described^7^. Total RNA was extracted, quality assessed, and sequenced on an Illumina NovaSeq platform. Sequencing reads were aligned to the Ensembl reference genome, and differential expression and pathway enrichment analyses were subsequently performed.

### Human Dorsal Root Ganglia Recovery, Culture, and Sequencing

Human DRGs were recovered from organ donors through the Southwest Transplant Alliance and cultured as previously described^26, 27^. A total of 8 DRGs from 3 donors were used (Supplemental Table 1). DRGs were dissociated, cultured under standard neuronal conditions, and treated with leptin (10 ng/mL) or vehicle control for 24 hours prior to RNA extraction. Detailed culture conditions, RNA extraction, and sequencing are provided in the Supplemental Methods.

### Human Infrapatellar Fat Pad and Synovium Single-Cell Integration

Secondary analysis of publicly available single-cell and single-nucleus RNA sequencing datasets from human infrapatellar fat pad and synovium^28, 29^ were integrated and analyzed using Seurat and Harmony^30–34^. Cell populations were identified using canonical marker genes, and leptin and leptin receptor expression patterns were assessed across cell types. Additional computational and analytical details are provided in the Supplemental Methods.

### Statistical Analysis

Statistical analyses were performed using GraphPad Prism (Version 10.6.1). Data are presented as mean with 95% confidence intervals (CI). Comparisons between groups were conducted using one-way or two-way analysis of variance (ANOVA), as appropriate and noted specifically in the figure legends, followed by Sidak’s or Tukey’s post hoc multiple comparisons test. Assumptions of normality and homogeneity of variance were assessed prior to parametric testing. Outliers were identified using Grubb’s test and excluded prior to analysis. A priori p-value was defined as 0.05.

## Results

### Chondrocyte-Specific Leptin Receptor Deletion Does Not Drive Osteoarthritis Pathology or Pain

To isolate the contribution of chondrocyte-intrinsic leptin signaling to joint damage and pain, we generated chondrocyte-specific leptin receptor knockout mice (LepR cKO) and subjected them to DMM. Deletion of the leptin receptor did not protect male or female mice from cartilage degeneration or osteophyte formation (Fig. 1A-C). However, female mice were protected from synovitis with DMM, relative to floxed controls, whereas males were not (Fig. 1D, E). Tamoxifen treatment alone did not affect behavioral outcomes (Supplemental Fig. 1). Similar pain-related behaviors were detected across groups among male and female mice (Fig. 1F). These findings demonstrate that leptin signaling within chondrocytes is neither required for OA-associated joint damage nor for the development of pain following joint injury. Together, these data indicate that chondrocyte-intrinsic leptin signaling does not account for OA-associated pain, prompting further investigation of extra-articular tissues and mechanisms underlying leptin-dependent effects in OA.

**Figure 1.**
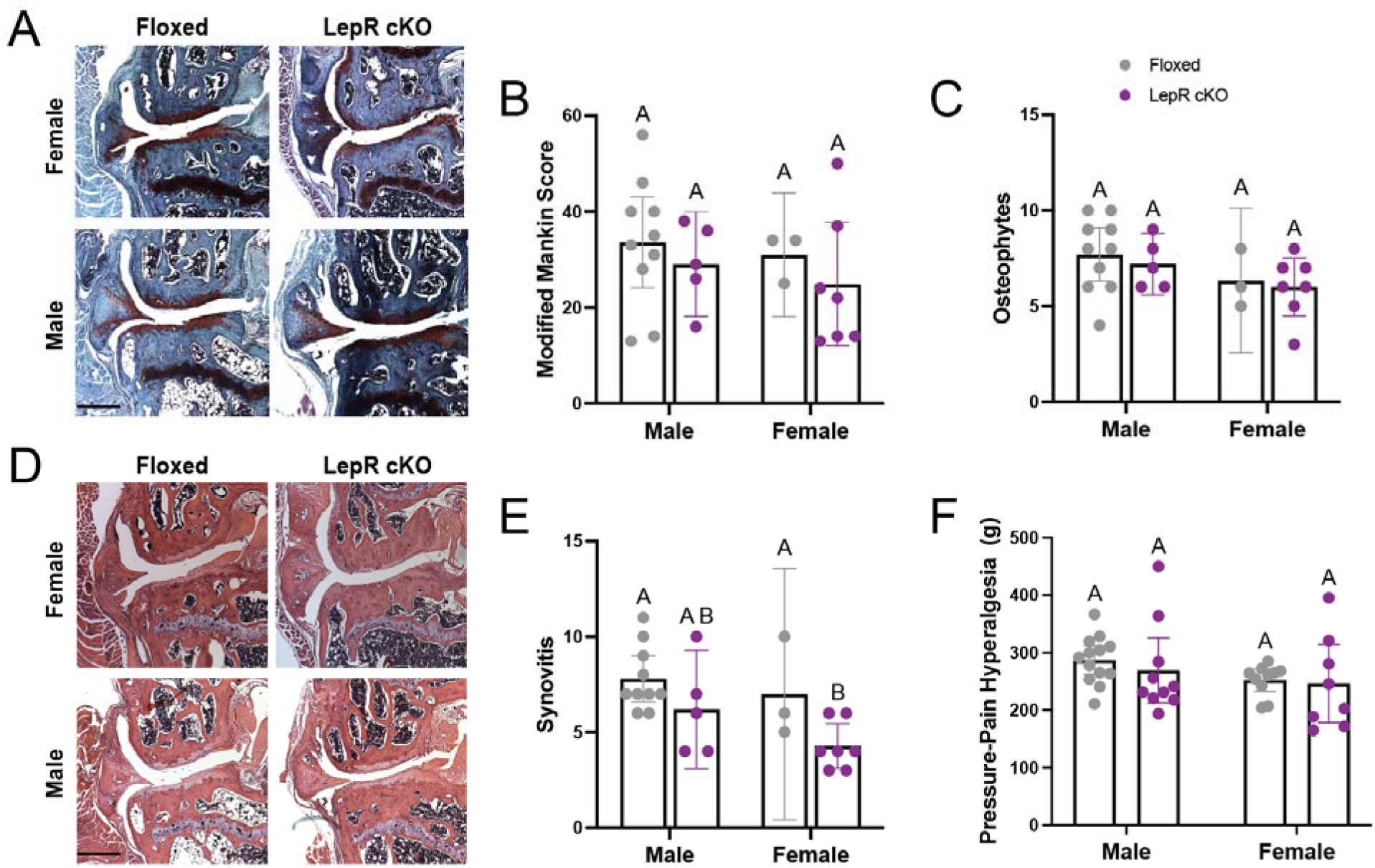
Chondrocyte specific deletion the leptin receptor does not prevent OA or pain. (A) Representative Safranin O/Fast Green and (D) H&E staining of male and female floxed (gray) and Chondrocyte-specific leptin knockout (LepR cKO, purple) mice DMM knee joints. Scale bar indicates 400 µm. Histological scoring including (B) Modified Mankin, (C) Osteophyte, and (E) synovitis assessments. Knee pain was assessed by (F) SMALGO in male and female DMM limbs. n = 3-13 per group. Data were analyzed by two-way ANOVA with Tukey, Sidak, or Bonferroni post hoc tests and are shown as mean with 95% confidence intervals. Different letters indicate p < 0.05.

### Global Leptin Deficiency Dissociates Osteoarthritis Pain from Structural Joint Damage

To define the systemic contribution of leptin to joint disease and nociception, we next examined WT, heterozygous leptin (Ob/+), and leptin-deficient (Ob/Ob) mice following DMM. Ob/Ob mice exhibited a pronounced obesity phenotype, with significantly increased body fat compared to WT and Ob/+ animals in both males and females (Fig. 2A, B). Histological analysis demonstrated that DMM induced cartilage degeneration across genotypes and sexes, as reflected by increased Modified Mankin scores in injured limbs relative to contralateral controls (Fig. 2C-F). In both males and females, Ob/+ mice demonstrated the lowest degree of cartilage damage with DMM, suggesting that partial leptin deficiency confers modest but consistent structural protection following joint injury. In males, WT and Ob/Ob mice exhibited comparable cartilage degeneration (Fig. 2C), whereas in females, Ob/Ob mice developed greater cartilage damage than both WT and Ob/+ mice – revealing a sex-dependent exacerbation of cartilage damage under conditions of complete leptin deficiency (Fig. 2D). Similarly, osteophyte formation was highest in the injured limbs of both male and female Ob/Ob mice compared to WT and Ob/+ animals, while Ob/+ mice remained the least affected (Fig. 2G, H). Synovitis scores were elevated in Ob/Ob mice following DMM in both sexes, and injured limb scores exceeding both WT and Ob/+ animals (Fig. 2I, J). Ob/+ mice showed a significant DMM-induced response in both sexes, but the pattern and magnitude were sex-dependent. In males, Ob/+ synovitis was lower than WT in response to DMM, consistent with partial structural protection, whereas in females, Ob/+ synovitis exceeded WT levels. Together, these structural findings reveal a consistent and perhaps unexpected pattern: partial leptin deficiency is structurally protective across multiple joint outcomes with and without DMM, while complete leptin deficiency markedly exacerbates joint damage.

**Figure 2.**
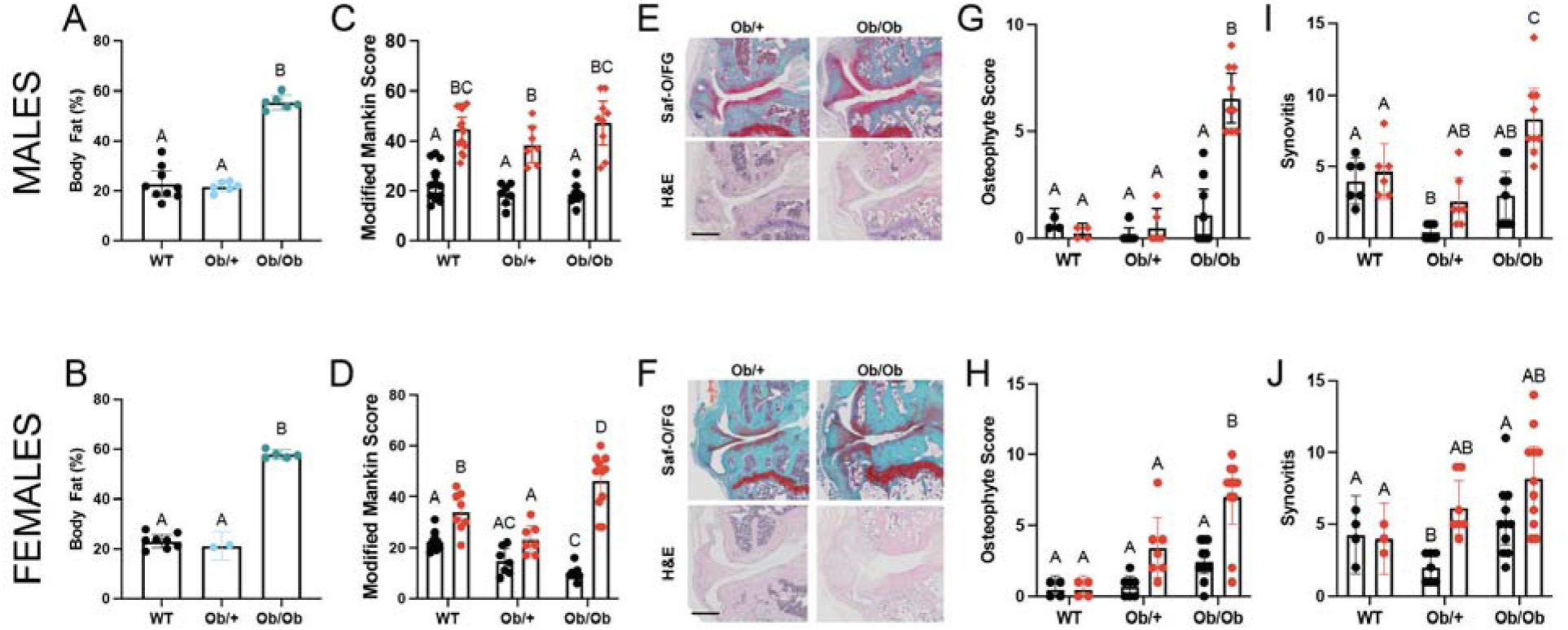
Global leptin deficiency exacerbates structural joint damage in males and females. Percent body fat in (A) male and (B) female WT (black), Ob/+ (blue), and Ob/Ob mice (green). Representative Safranin O/Fast Green and H&E staining of (E) male and (F) female Ob/+ and Ob/Ob knee joints. Scale bar indicates 400 µm. Histological scoring including (C,D) Modified Mankin, (G, H) osteophyte, (I, J) synovitis assessments where red indicates DMM limbs and black indicates contralateral limbs. n = 3-14 per group. Data were analyzed by one- or two-way ANOVA with Tukey, Sidak, or Bonferroni post hoc tests and are shown as mean with 95% confidence intervals. Different letters indicate p < 0.05.

In contrast, assessment of pain-related behaviors revealed a dissociation between structural joint outcomes and nociception, which did not demonstrate the overt sexual dimorphism that we observed in the structural phenotypes. Despite exhibiting the greatest degree of structural damage across all histological measures, Ob/Ob mice were protected from OA-associated pain in both sexes. Following DMM, WT and Ob/+ mice developed significant pressure-pain hyperalgesia in both sexes, as evidenced by a significant reduction in pain thresholds in the injured limb relative to contralateral controls. Notably, Ob/+ mice showed the greatest degree of hyperalgesia despite exhibiting the least structural damage (Fig. 3A, C). In contrast, Ob/Ob mice showed no significant difference between injured and contralateral limbs in either sex, indicating protection from DMM-induced hyperalgesia. Assessment of mechanical allodynia revealed a consistent pattern where Ob/Ob mice exhibited significantly elevated withdrawal thresholds, indicating less pain, compared to both WT and Ob/+ animals in both sexes. WT and Ob/+ thresholds were significantly different in males, but not in females (Fig. 3B, D). These data show that leptin is required for the development of OA pain, independent of structural joint damage.

**Figure 3.**
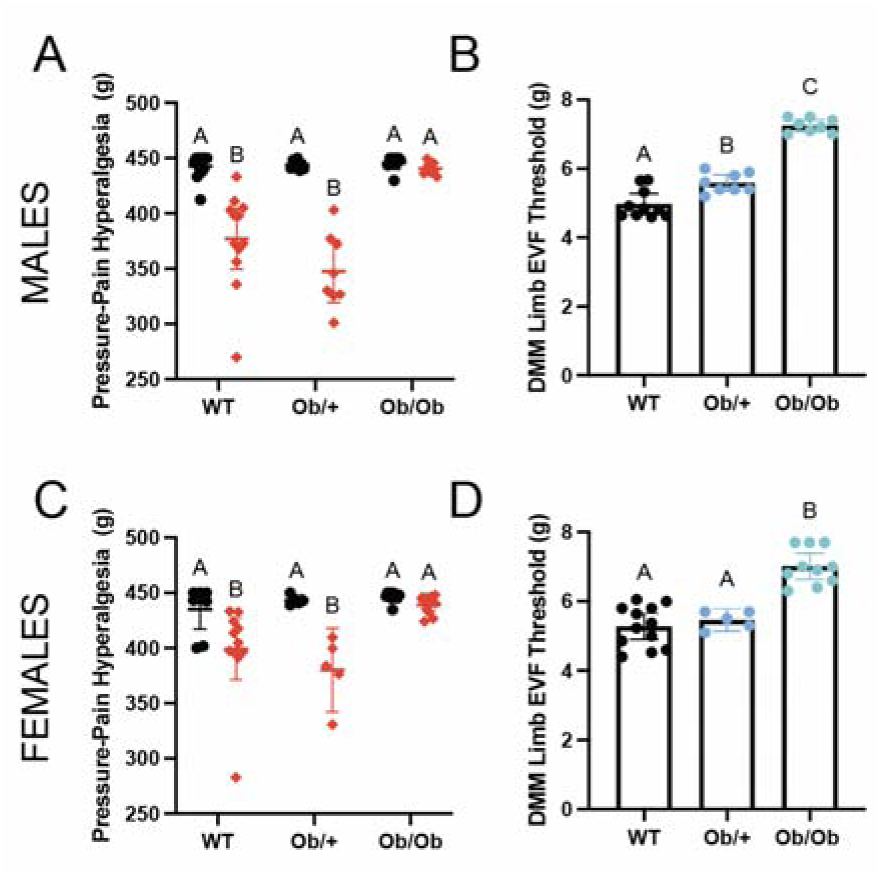
Leptin deficiency reduces OA-associated pain despite comparable structural damage. Knee pain was assessed by (A, C) SMALGO in male and female contralateral (black) and DMM limbs (red). (B, D) Electric Von Frey in DMM limbs of male and female WT (black), Ob/+ (blue), and Ob/Ob mice (green) was performed. n = 5-12 per group. Data were analyzed by one- or two-way ANOVA with Tukey, Sidak, or Bonferroni post hoc tests and are shown as mean with 95% confidence intervals. Different letters indicate p < 0.05.

To understand the impact of loss of leptin on the systemic inflammatory profiles, we profiled the serum immune landscape of these mice. As expected, Ob/Ob mice demonstrated increased circulating cytokines across genotypes in both sexes. In males, Ob/+ and Ob/Ob mice exhibited elevated levels of multiple pro-inflammatory mediators compared to WT controls, including IL-1α, IL-1β, IL-6, TNF-α, and MCP-1. Surprisingly, several cytokines were highest in Ob/+ mice, indicating a robust inflammatory state despite only partial leptin deficiency (Fig. 4A-F). In females, cytokine profiles were more variable across genotypes. While IL-1α was increased in Ob/+ and Ob/Ob mice relative to WT, other cytokines showed distinct patterns, with IL-6 and IL-10 highest in Ob/Ob mice and IL-1β, TNF-α, and MCP-1 highest in WT mice (Fig. 4G-L). Despite these sex-specific differences in circulating pro-inflammatory cytokines, which has been associated with OA pain before^35, 36^, leptin-deficient mice in both sexes remain protected from OA-associated pain. These findings indicate that reduced pain in leptin-deficient mice is not explained by cytokine profiles alone and instead support a role for leptin-dependent mechanisms within the joint or sensory nervous system in driving nociceptive sensitization.

**Figure 4.**
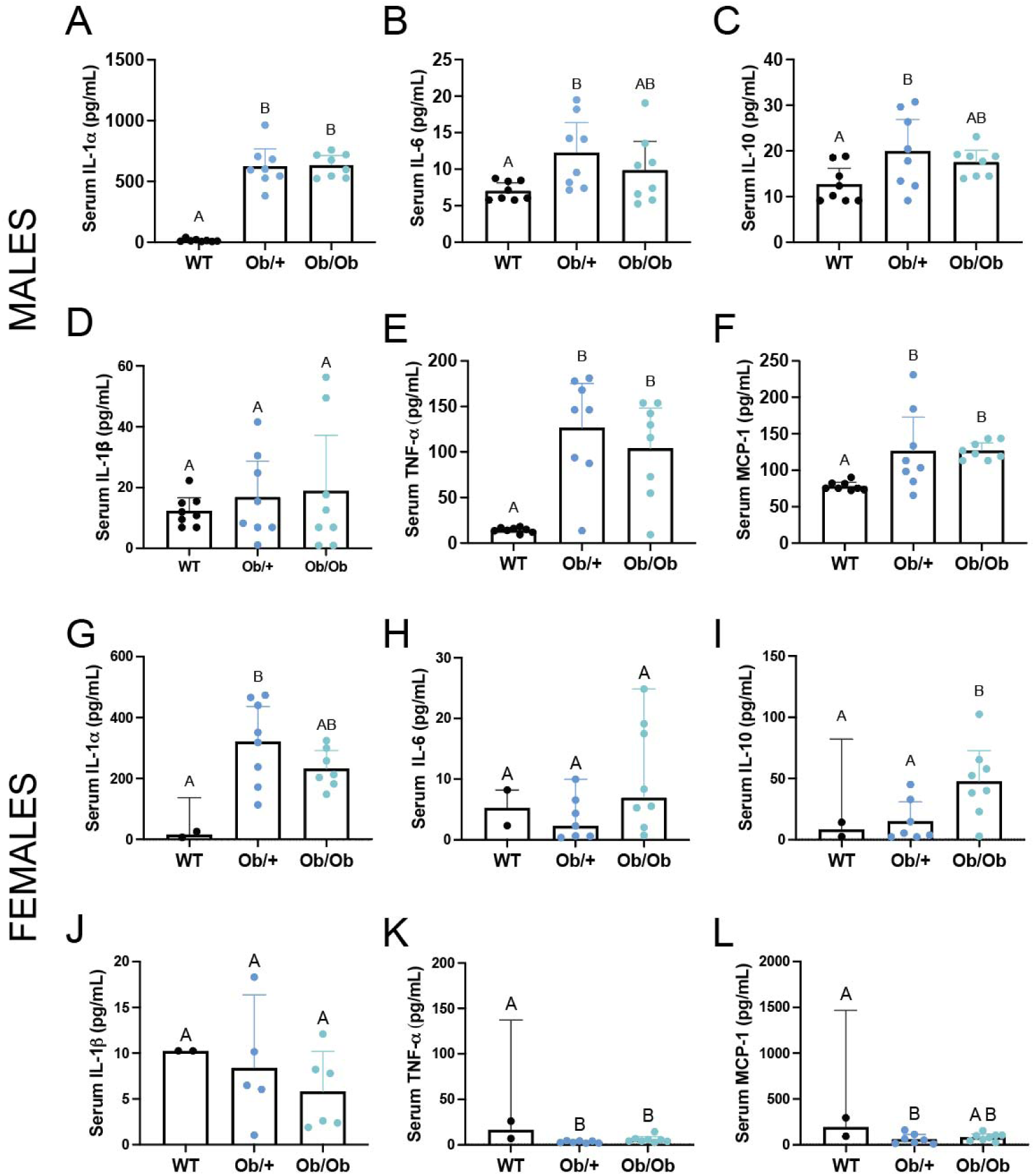
Leptin drives joint and systemic inflammation in OA. Serum levels of IL-1α, IL-6, IL-10, IL-1β, TNF-α, and MCP-1 were measured in (A-F) male and (G-L) female WT (black), Ob/+ (blue), and Ob/Ob mice (green). n = 2-8 per group. Data were analyzed by one-way ANOVA with Tukey, Sidak, or Bonferroni post hoc tests. Error bars represent the upper bound of the 95% confidence interval; values extending beyond the plotted range are truncated for visualization. Different letters indicate p < 0.05.

### Leptin Deficiency Alters Transcriptional Programs in DRG Neurons Innervating the Knee

Given that cartilage-specific leptin signaling did not account for OA-associated pain, we next investigated whether leptin influences transcriptional programs in DRGs that innervate the knee. To address this, we performed bulk RNA sequencing of L3-L5 DRGs from Ob/+ and Ob/Ob mice following DMM. Direct comparison of DRGs from Ob/+ and Ob/Ob mice following DMM revealed distinct transcriptional states with divergent pain phenotypes. Relative to Ob/+, Ob/Ob DRGs showed reduced expression of lipid and eicosanoid metabolism pathways and diminished inflammatory and neuronal sensitization (*Grin2b, Crabp2, Itih3, Gfap*), alongside increased expression of cytoskeletal and structural remodeling programs (*Serpina3i, Serpina3h, Adamts8, Cidea, Fst*) (Fig. 5A, Supplemental Table 2). These transcriptional differences paralleled pain outcomes, suggesting that leptin deficiency suppresses a key lipid-mediated inflammatory program in sensory neurons while promoting an alternative remodeling state that does not fully support nociceptive sensitization.

**Figure 5.**
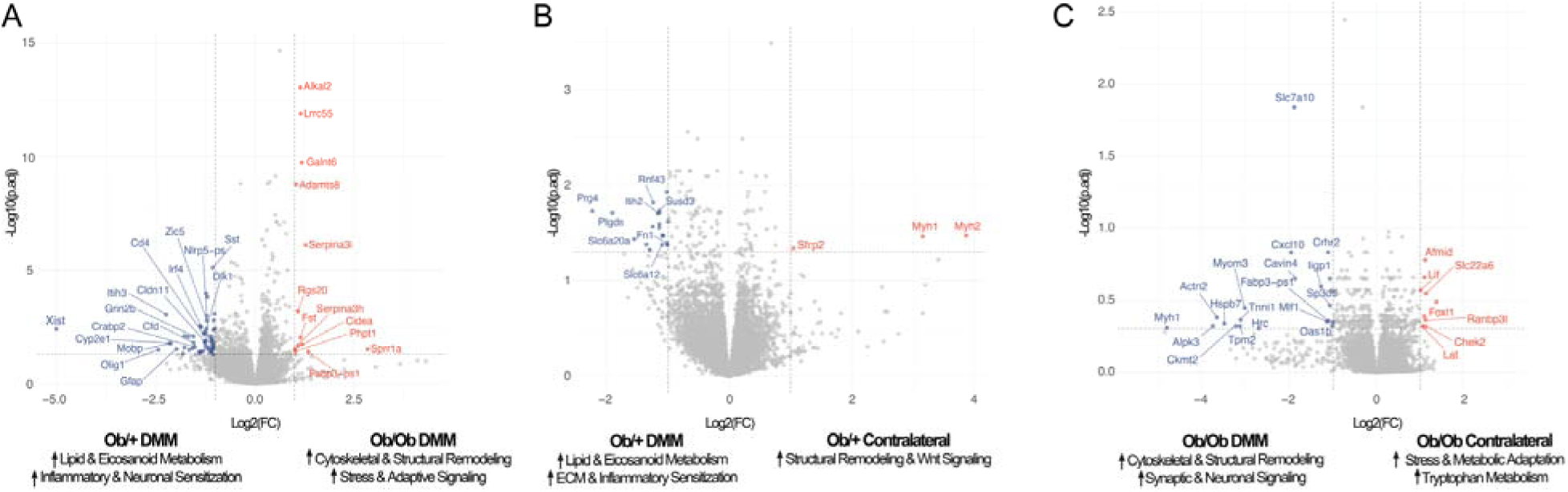
DRG neuron transcriptomic profiles reveal distinct leptin-dependent transcriptional programs associated with nociceptive sensitization. Volcano plots depicting differential gene expression in L3–L5 DRGs following destabilization of the medial meniscus (DMM). (A) Direct comparison of Ob/Ob vs Ob/+ DMM DRGs highlights increased lipid-mediated inflammatory signaling among Ob/+ DMM DRGs compared to increased cytoskeletal and structural remodeling pathways in Ob/Ob DRGs. (B) Ob/+ DMM vs contralateral DRGs show enrichment of lipid and eicosanoid metabolism, and extracellular matrix (ECM) and inflammatory pathways. (C) Ob/Ob DMM vs contralateral DRGs demonstrate enrichment of cytoskeletal and synaptic remodeling–associated programs. Differentially expressed genes are plotted as log2 Fold Change (Log2FC) versus -Log10(p-value), with selected biological themes annotated. Red – upregulated genes. Navy – downregulated genes. Grey – not significant.

Within-genotype comparisons further support these findings. In Ob/+ mice, DMM induced a transcriptional program consistent with nociceptor sensitization. Differential expression analysis revealed enrichment of pathway related to lipid and eicosanoid metabolism (Ptgds), inflammatory and extracellular matrix-associated signaling (*Fn1, Itih2*), and genes previously associated with pain and implicated in OA^37–41^ (*Fn1, Prg4, Itih2*) (Fig. 5B, Supplemental Table 3). In comparison, structural remodeling (*Myh1, Myh2*) and Wnt signaling (*Sfrp2*) were enriched in Ob/+ contralateral DRGs. In contrast, rather than engaging lipid-associated inflammatory pathways, Ob/Ob DRGs showed enrichment of cytoskeletal and structural remodeling (*Actn2, Myl1, Tpm2, Cavin4*), as well as pathways related to synaptic and neuronal signaling (*Hspb7, Slc7a10*) (Fig. 5C, Supplemental Table 4). This transcriptional profile is consistent with altered neuronal structure and signaling but notably lacks the lipid-inflammatory signature associated with nociceptor sensitization. Moreover, these findings align with the attenuated pain response observed in Ob/Ob mice despite more severe joint damage. In comparison, stress and metabolic adaptation (*Ranbp3l, Slc22a6, Lif*) and tryptophan metabolism (*Afmid*) were enriched in Ob/Ob contralateral DRGs.

Together, these findings indicate that leptin is associated with distinct transcriptional states in L3–L5 DRGs that innervate the knee following joint injury. In the presence of leptin, sensory neurons exhibit enrichment of lipid-mediated inflammatory pathways that are consistent with nociceptor sensitization, whereas in its absence, DRGs display a transcriptional profile characterized by cytoskeletal remodeling and altered neuronal signaling. While these data are derived from a single post-injury timepoint and do not directly assess neuronal function, the observed transcriptional differences parallel behavioral outcomes, suggesting that leptin-dependent changes in DRG gene expression may contribute to the dissociation between structural joint damage and pain observed in leptin-deficient mice.

### Leptin Treatment Reprograms Transcriptional State in Human Dorsal Root Ganglia Neurons

To explore the transcriptional response of human DRGs to leptin stimulation, we cultured human DRGs from organ donors and treated them with leptin 24 hours prior to RNA extraction. Bulk RNA sequencing and differential gene expression analysis revealed a total of 110 upregulated and 132 downregulated genes in response to leptin treatment. (Fig. 6A). Gene ontology analysis revealed enrichment of cell cycle progression driven by increases in *MKI67*, *CDK1*, and *TOP2A* and neuron projection due to increases in *CNTN2*, *KIF5A*, *EPHA3*, and *SCN10A* (Supplemental Table 5). Leptin treatment also altered the inflammatory state due to downregulation of *IL1B*, *IL6*, and *CCL4*. To further explore alterations in transcriptional programming induced by leptin treatment, GSEA analysis was employed, revealing that leptin treatment enriched neuronal associated programs including nociceptor excitability and ion channels, protein-protein interactions at synapses, and neurexins and neuroligins (Fig. 6B). The vehicle treated group again show enrichment of inflammatory pathways including TNF-α signaling via NF-κB, inflammatory response, and interferon response (Fig. 6B), suggesting that leptin influences inflammatory signaling within sensory neurons, potentially shifting the balance of immune–neuronal communication. Plotting paired leading-edge genes scores for the change in nociceptor excitability and ion channels and TNF-α signaling via NF-κB across each biological sample demonstrates similar transcriptional directionality across donors following leptin treatment (Fig. 6C, D).

**Figure 6.**
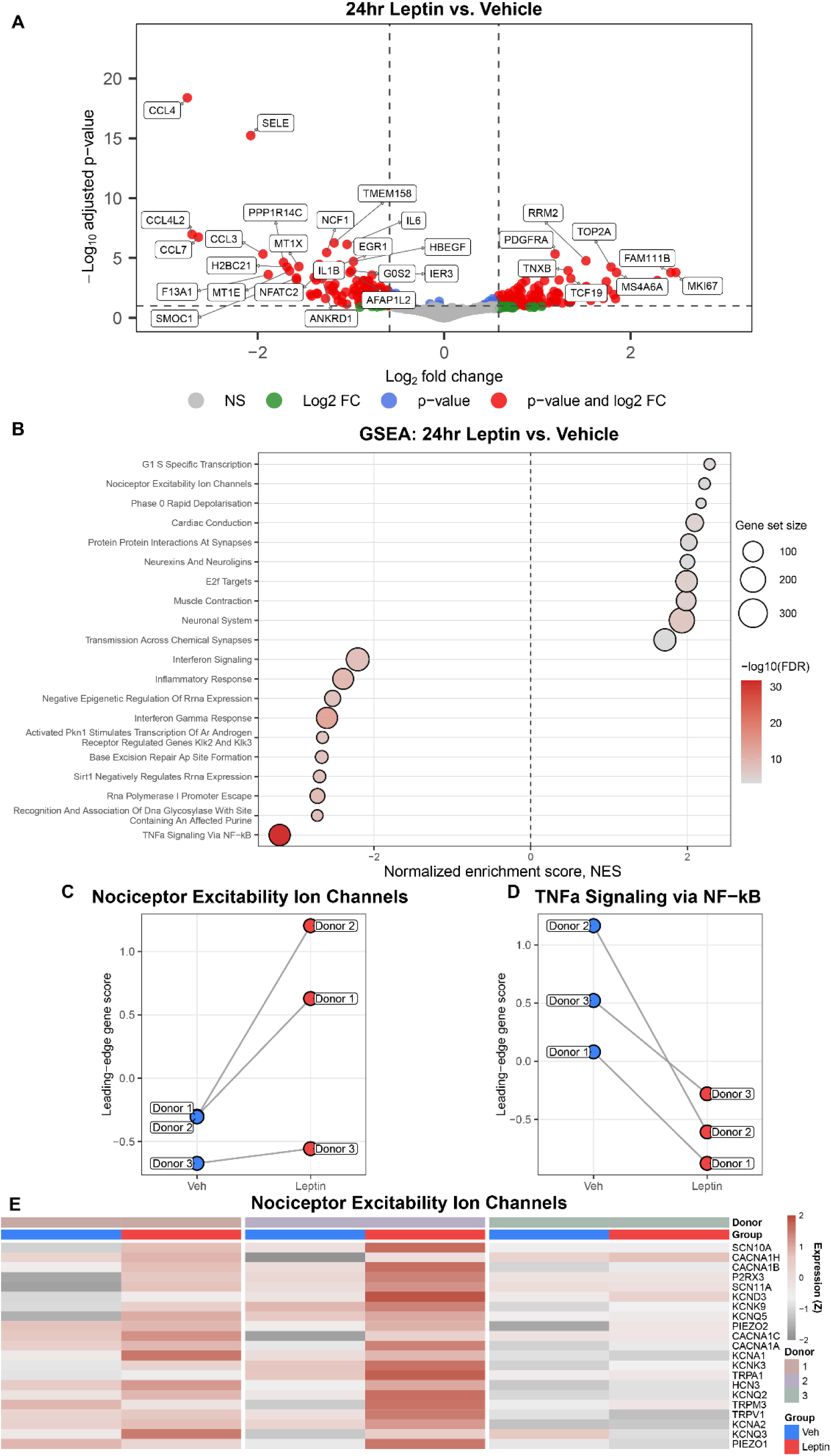
Leptin directly modulates transcriptional programs in human DRGs. (A) Volcano plot of protein coding differentially expressed genes identified via bulk RNA sequencing between leptin and vehicle treated cultured human DRG cultures. There were 110 upregulated genes and 132 downregulated genes. (B) Top 10 GSEA pathways ranked by adjusted p-value identified as enriched in leptin or vehicle treated cultures. Positive NES score denotes enriched in leptin treatment and negative score indicated enriched in vehicle treated cells. (C-D) Leading-edge gene scores for nociceptor excitability ion channels and TNFα signaling via NF-κB across biological replicates. Lines denote matched cultures from the same donor DRG. (E) Heatmap of VST gene expression of leading-edge genes from the nociceptor excitability ion channel pathway across each donor (n = 3) and treatment.

Building off this evidence that leptin plays a discordant role in development of pain and joint degeneration, we examined leptin’s effects on nociceptor associated transcriptional programming. Focusing on the leading-edge genes responsible for enrichment of nociceptor excitability and ion channels identified several known nociceptor enriched genes thought to play a direct role in pain processing and were increased including *SCN10A*, *P2RX3*, *PIEZO2*, and *TRPA1* (Fig. 6E). Together, this suggests that 24-hour leptin treatment shifts the transcriptome of cultured DRG cells toward increased cell cycle and proliferation, with enrichment of neuronal and nociceptor associated transcriptional programs, and decreases in inflammatory and cytokine signaling. Collectively, these data support a model in which leptin acts on sensory neurons, rather than chondrocytes, to influence the translation of joint injury into pain, providing a mechanistic explanation for the dissociation between structural joint pathology and nociception observed in leptin-deficient mice.

### Human Infrapatellar Fat Pad and Synovium Show Adipocyte-Enriched Leptin and Broad Leptin Receptor Expression

To provide human tissue context for the proposed adipose-sensory neuron axis, publicly available single-cell and single-nucleus RNA sequencing datasets from human infrapatellar fat pad and synovium were integrated and analyzed^28, 29^. Harmony integration identified and resolved major cell types including adipocytes, fibroblasts, endothelial cells, smooth muscle cells, macrophages, T/NK cells, B/plasma cells, and mast cells (Fig. 7A-C).

**Figure 7.**
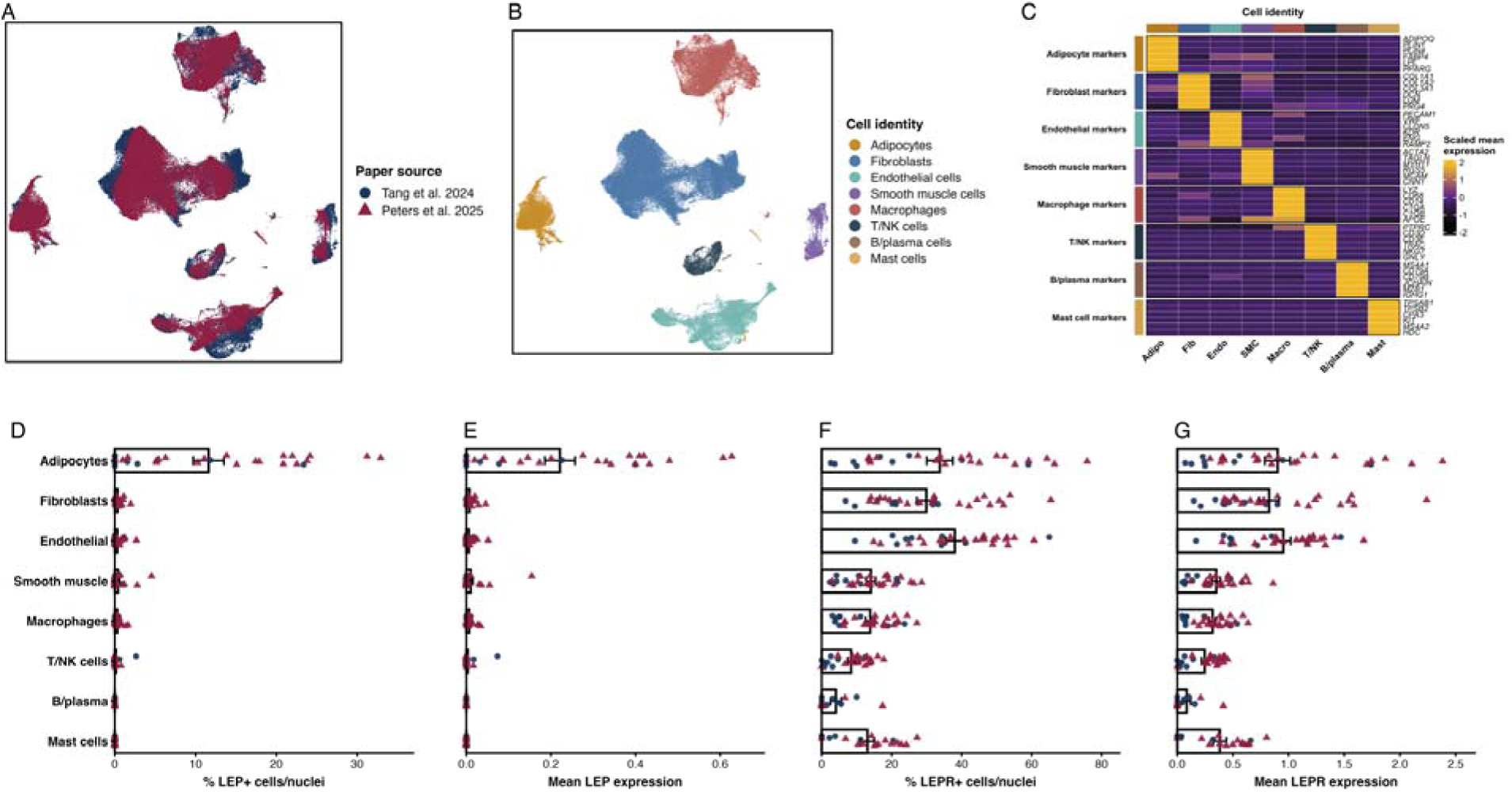
Integrated Lep and LepR expression in human infrapatellar fat pad and synovium. (A) UMAP colored by data source showing 140,842 cells/nuclei from 30 patient samples. (B) UMAP colored by cell type after labeling Harmony clusters with canonical marker genes. (C) Marker-gene heatmap showing scaled mean expression of canonical lineage markers across cell types. (D, E) Percentage of Lep-positive cells/nuclei and mean Lep expression per patient sample and cell type. (F, G) Percentage of LepR-positive cells/nuclei and mean LepR expression per patient sample and cell type. Data are shown as patient-level points overlaid on boxplot summaries. Lep expression is enriched in adipocytes, whereas LepR expression is detected across adipocyte, stromal, vascular, and immune cell types.

Leptin (*Lep*) expression was highly enriched in adipocytes, measured by the percentage of Lep-positive and LepR-positive cells/nuclei, relative to all other cell populations thus identifying adipocytes as the predominant local source of leptin within the infrapatellar fat pad and synovium (Fig. 7D, E, Supplemental Table 6). In contrast, leptin receptor (LepR) expression was broadly distributed across multiple cell types, including endothelial cells, fibroblasts, adipocytes, smooth muscle cells, macrophages, and mast cells (Fig. 7F, G, Supplemental Table 6). Collectively, these data support a model in which adipocytes inside and outside of the joint are the main local Lep-producing cell type, while LepR is expressed by multiple cell types that could respond to leptin inside and outside of the joint. This analysis provides human joint tissue context relevant to the study’s broader findings, suggesting that leptin-dependent pain signaling may involve cells outside cartilage, but within and outside the joint. Although this analysis does not demonstrate leptin signaling activity or pain mechanisms directly, the expression pattern is consistent with adipocyte-derived leptin acting locally on stromal, vascular, immune, and adipocyte cell types in the joint environment that warrants decoding in future studies.

## Discussion

The dissociation between structural joint damage and pain is one of the most clinically challenging and mechanistically unresolved features of OA, particularly in obesity. In the present study, constitutive leptin deficiency uncoupled nociception from structural pathology, revealing that obesity-associated OA pain is not solely determined by cartilage degeneration or synovitis. Despite exhibiting substantial joint pathology following DMM, leptin-deficient mice (Ob/Ob) were protected from hyperalgesia in both sexes, whereas heterozygous mice (Ob/+) developed robust pain despite comparatively less severe pathology. Together, these findings identify leptin as an important regulator of OA-associated nociceptive sensitization.

A key question raised by these findings is how leptin influences sensory neuron state at the molecular level. Transcriptomic profiling of knee-innervating DRGs identified enrichment of lipid and eicosanoid metabolism alongside inflammatory and nociceptor-associated signaling in the presence of leptin. Lipid mediators, including prostaglandins and arachidonic acid derivatives, are well-established drivers of peripheral sensitization that enhance sensory neuron excitability through receptor-mediated signaling pathways^42–46^. Leptin has also been shown to induce eicosanoid production in immune and joint-associated cells^15, 47, 48^. In contrast, DRGs from leptin-deficient mice failed to engage this transcriptional program and instead exhibited enrichment of cytoskeletal remodeling and structural neuronal pathways. This divergence provides a transcriptional explanation for the structure-pain dissociation: leptin determines whether sensory neurons enter a pain-permissive lipid-inflammatory state following injury.

Human DRG experiments extended these findings and provided translational evidence that leptin directly alters sensory neuron transcriptional states. Leptin stimulation enriched pathways associated with neuronal projection, synaptic organization, and nociceptor excitability, alongside increased expression of genes linked to sensory neuron function and pain processing, including *SCN10A, P2RX3, PIEZO2*, and *TRPA1.* In parallel, leptin suppressed TNF-α/NF-κB-associated signaling and reduced expression of inflammatory mediators including *IL-1β, IL-6*, and *CCL4*, suggesting that leptin reshapes inflammatory signaling within DRG cells while promoting nociceptor-associated programs. Importantly, these transcriptional patterns aligned closely with the *in vivo* DRG findings observed following DMM, supporting a conserved role for leptin in modulating sensory neuron state.

The present findings also challenge the assumption that leptin influences OA pain primarily through cartilage-intrinsic mechanisms. Although leptin has been implicated in cartilage metabolism and OA progression, chondrocyte-specific deletion of LepR did not alter structural outcomes or nociceptive behaviors in either sex, indicating that leptin signaling within chondrocytes alone is insufficient to account for pain phenotypes. One possibility is that leptin’s effects in obesity-associated OA arise through coordinated signaling across multiple joint-associated tissues rather than cartilage alone. Supporting this, analysis of human infrapatellar fat pad and synovium demonstrated adipocyte-enriched leptin expression alongside broad LepR distribution across stromal, vascular, immune, and adipocyte populations, suggesting that leptin signaling within the OA joint environment is multicellular and may involve coordinated interactions between adipose tissue and the sensory nervous system.

In obesity, where circulating leptin levels are chronically elevated, DRG neurons exhibited transcriptional programs enriched for lipid-mediated inflammatory and nociceptor-associated signaling pathways consistent with peripheral sensitization. In contrast, leptin deficiency was associated with cytoskeletal remodeling and altered neuronal signaling alongside reduced nociceptive behaviors despite substantial joint pathology. These findings suggest that while structural joint damage contributes to OA pain, the magnitude of nociceptive signaling may also depend on the transcriptional and functional state of sensory neurons. Leptin may therefore lower the threshold for DRG neuron activation in response to joint-derived signals, amplifying pain sensitivity in obesity-associated OA.

A notable strength of this study is the inclusion of both male and female mice across experimental cohorts. The dissociation between structural pathology and pain in leptin-deficient mice was observed in both sexes, suggesting that leptin-associated regulation of nociceptive signaling is not sex restricted. However, circulating cytokine profiles differed between males and females, consistent with prior work demonstrating sexually dimorphic leptin biology and adipose-associated inflammatory signaling^49^. Although the present findings do not resolve the mechanisms underlying these differences, leptin-dependent modulation of pain persisted despite divergent inflammatory profiles. Given that OA prevalence and pain severity are greater in females^50^, these findings underscore the importance of incorporating sex as a biological variable in studies of OA pain and obesity-associated signaling pathways while defining this adipose-pain crosstalk mechanism.

This study has limitations. Although the present findings support a role for sensory neuron–associated leptin signaling in OA pain, they do not directly establish that leptin receptor signaling within nociceptors is required in vivo. The observed phenotypes may reflect contributions from multiple LepR-expressing populations, and future studies using sensory neuron–specific LepR deletion models will be important to define cell type–specific mechanisms. In addition, the global leptin-deficient model introduces developmental and metabolic confounding variables, as Ob/Ob mice are obese and leptin-deficient from birth. Bulk RNA sequencing of whole DRGs also does not resolve cell type–specific transcriptional programs and likely reflects a composite of nociceptors, satellite glial cells, and infiltrating immune populations. Finally, although the human DRG experiments provide translational context, the *in vitro* culture system does not fully recapitulate the complexity of leptin signaling in the setting of joint injury and obesity.

These findings have important implications for the treatment of OA pain, which remains inadequately managed and disproportionately burdensome in obese individuals. Current therapeutic approaches largely target structural joint degeneration, yet the present findings suggest that structural pathology alone does not fully explain nociceptive outcomes. Interventions aimed at modulating sensory neuron signaling, including pathways downstream of leptin-associated lipid and inflammatory programs, may therefore provide an opportunity to reduce OA pain independent of structural disease modification. More broadly, these findings support the concept that obesity-associated OA pain may arise, in part, through adipose–sensory neuron signaling pathways that sensitize nociceptive circuits following joint injury.

## Acknowledgements

This study was supported in part by the Arthritis National Research Foundation, University of California Training for Research on Aging and Chronic Disease (AG049663), Shriners Hospitals for Children, National Institutes of Health grants including the Director’s New Innovator Award (DP2AG093209), R00AR078949, AG46927, AR072999, AR074992, P50 CA094056 (Molecular Imaging Center), R01NS111929, F32NS13563, K01AR079045, PRECISION Human Pain Network (RRID: SCIR_025458) part of the NIH HEAL Initiative under award U19NS130608 (awarded to T.J.P), Rheumatic Diseases Research Resource-based Center P30 AR073752, and NCI P30 CA091842 (Siteman Cancer Center Small Animal Cancer Imaging shared resource). Additional NIH support included funding from the National Institute of Neurological Disorders and Stroke through the PRECISION Human Pain Network (RRID:SCR_025458), part of the NIH HEAL initiative (https://heal.nih.gov/) under award number U19NS130608 to TJP. Additional support was provided by the Arthritis Foundation, the Nancy Taylor Foundation for Chronic Diseases, and the Philip and Sima Needleman Fellowship from the Washington University Center of Regenerative Medicine. We thank Dr. Martin Myers for providing LepR^fl/fl^ mice. A Pilot and Feasibility grant was awarded to K.H.C. from the Chicago Center for Musculoskeletal Pain at Rush University (P30 AR079206), which supported the bulk RNA-seq on the DRGs.

## Author Contributions

Conceptualization – HDW, KLL, BAA, AKO, JBL, DEO, TJP, FG, KHC. Methodology – HDW, AKO, JBL, DEO, TJP, FG, KHC. Investigation – HDW, EKT, KLL, BAA, MRM, REV, JBL, DEO, TJP, FG, KHC. Visualization – HDW, EKT, KLL, BAA, MRM, REV, JBL. Funding acquisition – TJP, FG, KHC. Project administration – HDW, JBL, DEO, TJP, FG, KHC. Supervision – HDW, JBL, DEO, TJP, FG, KHC. Writing – original draft – HDW, EKT, JBL. Writing – review & editing – HDW, EKT, KLL, BAA, MRM, REV, AKO, JBL, DEO, TJP, FG, KHC.

## Supplemental Figures

**Supplemental Figure 1.**
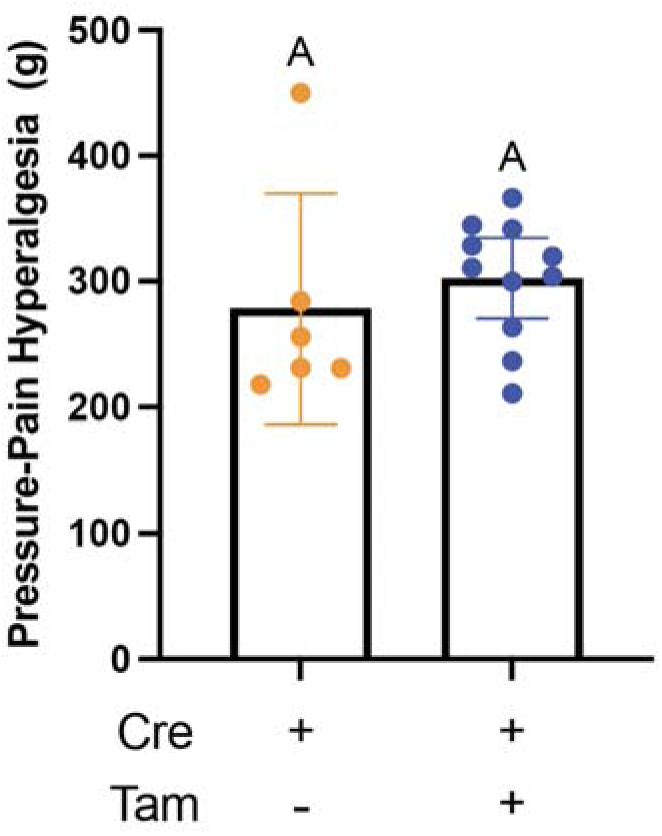
Tamoxifen treatment does not alter pain-related behavioral outcomes. Pressure pain threshold of DMM limbs were measured using SMALGO in Cre-positive LepR cKO mice with (blue) or without tamoxifen treatment (orange). Cre-positive LepR cKO mice. No differences in pressure-pain thresholds were observed between groups, indicating that tamoxifen treatment alone does not affect behavioral outcomes. n = 6-11 per group. Data were analyzed by Welch’s t-test and are shown as mean with with 95% confidence intervals. Different letters indicate p < 0.05.

## Supplemental Tables

**Supplemental Table 1.** Organ donor demographic data.

**Supplemental Table 2.** Transcriptional pathway analysis of differentially regulated genes between knee-innervating DRGs of DMM limbs of Ob/Ob (up) and Ob/+ (down) limbs.

**Supplemental Table 3.** Transcriptional pathway analysis of differentially regulated genes between knee-innervating DRGs of Ob/+ contralateral (up) and DMM (down) limbs.

**Supplemental Table 4.** Transcriptional pathway analysis of differentially regulated genes between knee-innervating DRGs of Ob/Ob contralateral (up) and DMM (down) limbs.

**Supplemental Table 5.** Gene Ontology results – including biological process, cellular component, and molecular function – of human DRGs in response to leptin stimulation.

**Supplemental Table 6.** Exact patient-level Lep and LepR expression summary by cell type. Values are medians with observed ranges across patient-level cell-type summaries from Fig. 7D-G. Patient sample/cell-type combinations with fewer than 10 cells/nuclei were excluded from the summary, matching the plotted analysis.

## Supplemental Methods

### Human Dorsal Root Ganglia Recovery and Culture

In collaboration with the Southwest Transplant Alliance, human DRGs were recovered from organ donors and cultured as previously described^1, 2^. All human tissue procurement procedures are approved by the University of Texas at Dallas IRB under protocol Legacy-MR-15-237. Standard procedures and policies for donor screening and consent, and tissue recovery are handled by the Southwest Transplant Alliance and are authorized by the United Network for Organ Sharing and the US Centers for Disease Control. All use and circulation of any donor medical information strictly follows HIPAA regulation. A total of 8 DRGs were used from 3 organ donors. Donor demographic information is provided in Supplemental Table 1.

Briefly, DRGs were surgically recovered within 4 hours of cross clamp and placed in a CSF or Hibernate A (∼4hrs) prior to dissociation as previously described^3^. The DRG bulbs were then trimmed of excess connective tissue and fat, diced into 3mm X 3mm sections and placed in 5 mL of HBSS without calcium and magnesium (Thermo Scientific, 14170-112) containing 1mg/mL of Stemxyme 1 (Worthington Biochemical, LS004106), 0.1 mg/mL of DNAse 1 (Worthington Biochemical, LS002139) and 10ng/mL of human β-NGF (R&D Systems, 256-GF). The tubes were placed in a shaking water bath set at 37°C and were triturated with glass, fire polished Pasteur pipettes every hour until the DRG sections were dissolved (4-5hrs). The following cell suspension was filtered through a 100µM mesh strainer (Corning, 431752) and then layered over 3 mL of a 10% BSA solution in HBSS in a 15 mL tube. The tubes were centrifuged for 5 minutes at 900g at room temperature. The supernatant was aspirated and the cellular pellet was resuspended in pre-warmed DRG media (BrainPhys^®^ media (Stemcell technologies, 05790), containing 1% penicillin/streptomycin, 2% NeuroCult SM1, 1% Glutamax, 1% N-2, 10ng/mL β-NGF (R&D Systems, 256-GF-100), 2% HyClone™ Fetal Bovine Serum (Thermo Fisher Scientific, SH3008803IR), and 0.1% 5-Fluoro-2′-deoxyuridine (FRDU, Sigma-Aldrich, F0503). The cells were then plated on 6 well glass bottom plates (P06-1.5H-N) pre-coated with 0.1mg/mL of poly-D-lysine (Sigma-Aldrich, P7405-5MG). The cultures were placed in an incubator set at 37°C and 5% CO2 for 3 hours to allow for neuronal adherence. Following adherence, wells were flooded with prewarmed media, and half media changes were performed every other day. Leptin treatments (10ng/mL; R&D Systems, 398-LP) or it’s vehicle (20 mM Tris-HCl, 0.1% final volume) were delivered to cell cultures on DIV 4 for a total treatment time of 24 hours.

### RNA Extraction

Following leptin treatment, RNA was extracted from cultured DRGs as previously described using Qiagen micro kit following the manufacturer’s recommended protocol (Qiagen, 74034)^2^. The only alteration in the protocol was the addition of an extra on-column DNAse clean up step (Qiagen, 79256). RNA samples were stored at -80°C, prior to subsequent analysis. All RNA samples had RIN values >8.0.

### Human DRG Bulk RNA Sequencing

RNA sequencing was performed using a NextSeq2000 for paired-end, mRNA sequencing at the University of Texas at Dallas. Raw FASTQ files were quality-controlled with FastQC v0.11.0 and analyzed for Phred scores, per-base sequence, and duplication levels. Sequencing reads were soft-clipped (12 bases per read) to remove adapters and low-quality bases, then aligned and sorted using STAR v2.7.6 with the GRCh38 human reference genome (GENCODE release 38, primary assembly). Deduplication was performed using Sambamba v0.8.2 and ENCODE blacklist genes were discarded using bedtools (v2.30.0). Normalized transcripts per million (TPM) for each gene was obtained using StringTie v2.2.1. The Rsubread (v2.14.2) featureCounts function was used to calculate raw counts for all genes. Differential gene expression analysis was conducted using DESeq2 to explore for differences between leptin and vehicle treated samples while controlling for Donor. Genes were considered differentially expressed if *P_adj_*< 0.1 and log_2_ fold-change (FC) > |±0.585|. Upregulated and downregulated genes were analyzed for Gene Ontology (GO) biological process, cellular component, and molecular function using EnrichR. A gene set enrichment analysis (GSEA) was performed using Hallmark and Reactome genes sets from molecular signature database along with custom DRG gene sets designed to test specific neuronal and nociceptor excitability transcriptional programs. Enrichment results were analyzed using normalized enrichment scores (NES) with false discovery rate adjusted p-values. Positive NES values indicated enrichment in leptin treated conditions while negative values signify enrichment in vehicle treated cells.

### Human Infrapatellar Fat Pad and Synovium Single-Cell Integration

Publicly available single-cell and single-nucleus RNA sequencing datasets from human infrapatellar fat pad and synovium were obtained from Tang et al. and Peters et al^4, 5^. The integrated dataset included 140,842 cells/nuclei from 30 patient samples, including 9 Tang samples and 21 Peters samples. Analyses were performed in R using Seurat for single-cell object handling, normalization, expression summarization, and dimensionality reduction, and Harmony to reduce dataset/source effects^6, 7^. The Harmony-corrected data were visualized by UMAP^8^. Data wrangling and patient-level summaries were performed using tidyverse packages including dplyr and tidyr, and figures were generated using ggplot2, ComplexHeatmap, and patchwork^9, 10^.

Clusters were labeled as adipocytes, fibroblasts, endothelial cells, smooth muscle cells, macrophages, T/NK cells, B/plasma cells, or mast cells using canonical lineage marker genes. Cell labels were checked using canonical marker-gene expression. For each patient sample and cell type, Lep-positive and LepR-positive cells/nuclei were defined as cells/nuclei with detectable nonzero expression of the corresponding gene. The percentage of Lep-positive or LepR-positive cells/nuclei was calculated as the number of positive cells/nuclei divided by the total number of cells/nuclei in that patient sample and cell type. Mean Lep and LepR expression were also calculated for each patient sample and cell type. Patient-level values were plotted as points overlaid on boxplots. These analyses were descriptive; statistical comparisons were not performed for the patient-level Lep/LepR summary panels.

